# Exploring G-quadruplex structure in *PRCC-TFE3* fusion Oncogene: Plausible use as anti cancer therapy for translocation Renal cell carcinoma (tRCC)

**DOI:** 10.1101/2023.07.06.547934

**Authors:** Neha, Parimal Das

**Affiliations:** Centre for Genetic Disorders, Institute of Science, Banaras Hindu University, Varanasi- 221005. Uttar Pradesh, INDIA

**Keywords:** *PRCC-TFE3* Fusion gene, Papillary Renal Cell Carcinoma, Xp11.2 translocation, G-quadruplex

## Abstract

The *TFE3* fusion gene, byproduct of Xp11.2 translocation, is the diagnostic marker for translocation renal cell carcinoma (tRCC). Absence of any clinically recognized therapy for tRCC, pressing a need to create novel and efficient therapeutic approaches. Previous studies shown that stabilization of the G-quadruplex structure in oncogenes suppresses their expression machinery. To combat the oncogenesis caused by fusion genes, our objective is to locate and stabilize the G-quadruplex structure within the *PRCC-TFE3* fusion gene. Using the Quadruplex-forming G Rich Sequences (QGRS) mapper and the Non-B DNA motif search tool (nBMST) online server, we found putative G-quadruplex forming sequences (PQS) in the *PRCC-TFE3* fusion gene. Circular dichroism demonstrating a parallel G-quadruplex in the targeted sequence. Fluorescence and UV-vis spectroscopy results suggest that pyridostatin binds to this newly discovered G-quadruplex. The PCR stop assay, as well as transcriptional or translational inhibition by PQS, revealed that stable G-quadruplex formation affects biological processes. Confocal microscopy of HEK293T cells transfected with the fusion transcript confirmed G-quadruplexes formation in cell. This investigation may shed light on G-quadruplex’s functions in fusion genes and may help in the development of therapies specifically targeted against fusion oncogenes, which would enhance the capability of current tRCC therapy approach.

## 1. Introduction

Translocation Renal Cell Carcinoma (tRCC) is a well-recognized sub-type of renal cell carcinoma categorized under a distinct group by the WHO (World Health Organization) classification system in 2004[1]. The Xp11.2 translocation is mostly proclaimed tRCC, characterized by translocation in the TFE3 gene resulting in the *TFE3* fusion gene product. *TFE3* have numbers of reported fusion gene partners such as *CLTC* [2], *ASPL, NONO* [3], *PRCC* [4], *RCC17* [5], *SFPQ* [6], *RBM10* [7] etc. Many cancers have been reported to associate with Xp11.2 translocation viz. *SFPQ-TFE3, ASPL-TFE3, YAP-TFE3*, *PRCC-TFE3* fusion gene present in Perivascular Epithelioid Cell neoplasms (PECOMAs), Alveolar Soft Part Sarcoma (ASPS), Epithelioid Hemangio-Endothelioma (EHE), and Papillary Renal cell Carcinoma respectively [8–10]. TFE3 is IgH enhancer-binding transcription factor, a 576 aa protein containing a basic helix– loop–helix domain for DNA binding, an acidic activation domain at N-terminal and a proline-rich activation domain at C-terminal, and a leucine zipper region for dimerization. *PRCC* is a Proline-Rich Mitotic Checkpoint Control Factor gene on chromosome 1 encoding PRCC protein of 491 amino acids carrying proline, leucine, and glycine-rich N-terminal region. t(X;1)(p11.2;q21) translocation causes fusion of Proline-Glycine rich N-terminal of *PRCC* with C-terminal of TFE3 resulting in *PRCC-TFE3* fusion gene having enhanced transcriptional activation [4,11,12]. The *PRCC-TFE3* fusion gene is a more potent transactivator than wild *TFE3* which results in aberrant expression of *TFE3* targeted genes. Skalsky Y M. *et al*[12]. reported that only the *PRCC-TFE3* fusion enhanced the transcriptional activation from endogenous *TFE3* proposing that the *TFE3* fusions are not consistent with the enhancement of transcriptional activation [12–13]. A *PRCC-TFE3* fusion gene has confirmed two different breakpoints in *PRCC*. In one breakpoint 156 amino acids of the N-terminal of *PRCC* while in the other 393 amino acids of the N-terminal of *PRCC* fused to *TFE3* [11,13]. The t(X;1)(p11.2;q21.2) translocation is predominantly reported in papillary renal cell tumors more frequently in young onsets. In all three sub-type of papillary RCC, the t(X;1)(p11.2;q21.2) is found to be associated with loss of wild *TFE3* and generation of *PRCC-TFE3* [4]. These findings instigate that the *PRCC-TFE3* fusion gene is possibly responsible for cell transformation in papillary renal cell carcinoma and could be an ideal target for an anti-cancer approach for papillary renal cell carcinoma. In the present study we have tried to find out an alternative way to target the *PRCC- TFE3* fusion gene by G-quadruplex structure.

G-quadruplexes are a special non-B form of DNA secondary structures formed by guanine-rich sequences which get self-assembled through multiple cyclic Hoogsten hydrogen bonds into a four-stranded helical structure. Putative G-quadruplex forming sequences (PQS) are systematically distributed within human genomes. G-quadruplex forming sequences are found in functional domains of several genes viz. telomeric region of a eukaryotic chromosome, 5’-UTR of mRNA, the promoter region of several oncogenes, etc. whose stabilization down-regulate the expression of corresponding genes [13–14]. Several findings suggest that the G-quadruplex stabilization in target genes can down-regulates their expression since the G-quadruplex structure within the regulatory & promoter site hampers transcription and translation by creating blockage in further processing of polymerase enzyme [15–16]. Based on these information, it could be hypothesized that stabilizing a G-quadruplex structure within the *PRCC-TFE3* fusion gene, an oncogene in t(X;1)(p11.2;q21.2) mediated papillary renal cell carcinoma could be an efficacious attempt to suppress their expression that could be a potential therapeutic approach for fusion gene associated oncogenesis. We have reported the PQS within *PRCC-TFE3* fusion oncogene in our published review article [13], in the present study we investigate the possibility of folding this PQS into G-quadruplex *In vitro* and showed its consequence over fusion gene expression.

## 2. Materials and Methods

### 2.1 Materials and Reagents

Pyridostatin (PDS) were purchased from Sigma Aldrich. Dual Luciferase assay kit and PsiCHECK^TM^-2 vector were purchased from Promega. Oligonucleotides were synthesized and obtained from Eurofins Scientific. HEK293T (Human embryonic kidney) cell line and Xp11.2 translocation renal cell cancer cell line UOK 146 with *PRCC-TFE3* translocation were cultivated in 5% (v/v) CO2 in a humid incubator while being maintained in DMEM supplemented with 10% fetal bovine serum and antibiotic cocktail (100 U/ml penicillin, 100 g/ml streptomycin). T4- PNK (polynucleotide kinase) and NheI enzymes were purchased from New England Biolab (NEB).

### 2.2 Sample Preparation

All oligonucleotides were desalted and HPSF purified. The oligonucleotides listed below were utilized in this study.

i. *PRCC* (G4Q): 5’-GTTGGGGAGGGACTGGGATTGGGGTTG-3’ (sequence within the *PRCC* part of *PRCC-TFE3* fusion gene)
ii. Mutated *PRCC* (G4M): 5’-GTTGAGGAGAGACTGAGATTGAGGTTG-3’
iii. Human Telomeric (TelQ): 5’- AGGGTTAGGGTTAGGGTTAGGG-3’

In order to generate a stable G-quadruplex structure, the oligos were incubated at 95°C for 5 min (minute) in a solution of 10 mM Tris and 100 mM KCl pH 7.4. The samples were then gradually cooled at room temperature for overnight.

### 2.3 Circular Dichroism Spectroscopy

A quartz cuvette with a 2.0 mm path length was used to measure CD spectra in a JASCO J-1500 CD Spectrophotometer with a wavelength range of 220-320 nm. The instrument’s scanning speed was adjusted to 50 nm min^-1^, and the data interval time employed was 1 sec. Bandwidth was 1.0 nm with three continuous scans and a data pitch of 0.2 nm. The oligonucleotides tested had a 5 µM strand concentration. Each spectrum consisted of an average of three observations taken at 25°C. In addition, the sample’s spectra was captured and subtracted from the spectrum of the respective buffer. Using Origin 8.0 software, all spectra were baseline-corrected and examined.

### 2.4 UV Vis Spectroscopy

UV-vis absorption assessment is a feasible approach that can be used to examine structural changes and complex information [32]. Absorption spectra were measured on a UV-1800 UV/Visible Scanning Spectrophotometer with a 1 mm path-length quartz cell and were recorded in the 200-800 nm range at 25°C. With stepwise addition of folded PRCC G4Q and G4M oligo sample to a UV-cell containing 5 μM solutions PDS, UV-Vis absorption titrations were performed. The measurements were carried out in a 10 mM Tris solution with a pH of 7.4 and 100 mM KCl.

### 2.5 Steady State Fluorescence Spectroscopy

To monitor the binding of PDS to the *PRCC* G-quadruplex DNA structure, changes in fluorescence spectra of 5 µM of PDS was recorded by increasing the concentration of *PRCC* G4Q & G4M. Fluorescence spectra were captured using Perkin Elmer LS55 fluorescence spectrometer. All measurements were performed in quartz cuvette with a 1 mm path length. PDS was excited at 268 nm, and the emission spectra were observed between 500 and 600 nm. The excitation and emission bandwidths were 5 nm. A stock solution of oligonucleotides (G4Q and G4M) was added in increasing amounts (0-10 μM) to a 5 µM of PDS, and fluorescence emission was measured after 2-5 minutes following each addition.

### 2.6 Thermal Melting Study

A thermal melting study was conducted using a JASCO J-1500 CD Spectrophotometer that has a Peltier temperature controller. G-quadruplex samples were prepared as mentioned in the sample preparation section. 5 µM of the G-quadruplex sample was mixed with 5 µM PDS and incubated at room temperature for 1-2 min. Samples were heated from 30°C to 90°C at 1°C per min temperature gradient. Absorbance data were recorded at 295 nm, as previous data showed that the wavelength of 295 nm produced more accurate findings than 260 nm for examining G- quadruplexes [15].

### 2.7 PCR Stop Assay

In this study, *PRCC*-R (5’TCTCCATCCCCTCTCAACC’3) a partly complementary oligonucleotide that moderately hybridizes to the 3’ end of the G-repeat of the G4Q oligo bearing a G-quadruplex sequence or its mutant (G4M) oligos, was utilized. The experiments were performed according to Verma *et al*. 2018 [11], In brief, in 1X PCR buffer with 3 units of Taq DNA polymerase, 0.05 mmol/l dNTPs, and 20 pmol oligonucleotides with variable PDS concentrations (0,2,3,5,7,&10 µM) the chain extension reaction was carried out. PCR conditions: 94°C for 5 min, followed by 30 cycles of 94°C for 30 s, 55°C for 30 s, and 72°C for 30 s, with a final extension at 72°C for 10 min. The amplified PCR products were resolved on 3% Agarose gels.

### 2.8 Clone Preparation

The psiCHECK-2 plasmid (Promega) has a special NheI restriction site that was used to insert a synthetic DNA duplex encoding the G-rich region of the *PRCC-TFE3* fusion gene in the upstream of the renilla luciferase initiation codon to create the plasmids G4Q27 and G4M27 (mutated G-rich region of the *PRCC-TFE3* fusion gene). For this NheI restriction sequence, CTAG was incorporated at the 5’ end of each set of oligos. A DNA oligonucleotide sequence from the following list was used:

*PRCC* G4Q sense: 5’ CTAGGTTGGGGAGGGACTGGGATTGGGGTTG 3’

*PRCC* G4Q antisense: 5’ CTAGCAACCCCAATCCCAGTCCCTCCCCAAC 3’

*PRCC* Mut G4Q sense: 5’ CTAGGTTGAGGAGAGACTGAGATTGAGGTTG 3’

*PRCC* Mut G4Q antisense: 5’CTAGCAACCTCAATCTCAGTCTCTCCTCAAC3’

Each set of antisense and sense oligonucleotides was annealed at 5 µg concentration by heating at 95°C for 5 min with annealing buffer (100 mM potassium acetate, 30mM HEPES PH 7.5) followed by overnight gradual cooling at room temperature. Annealed duplex oligos were quantified. After that, the non-radioactive phosphorylation of the produced duplex oligos was done by using T4-PNK at 37°C for 30 min, followed by 65°C for 20 min. The psiCHECK-2 vector was digested with the NheI restriction enzyme (single digestion), followed by CIP (calf intestinal phosphatase) treatment to form sticky ends, which were then ligated with duplex oligos in a 1:1 ratio into the upstream of the renilla luciferase start codon of the psiCHECK-2 plasmid. The ligated product was then transformed into *E. coli* cells, and the plasmid was isolated using a miniprep. The extracted plasmid was sequenced in order to validate the creation of a clone (insert) and was subsequently used for transfection (Schematic presentation of reporter plasmid shown in Figure 10). Additionally, full-length *PRCC-TFE3* fusion transcript cDNA was cloned into the pcDNA3.1 expression vector to make a fusion construct plasmid. The constructs were sequence verified by DNA sequencing.

### 2.9 Dual Luciferase Assay

For the dual luciferase experiment, sub-confluent HEK293T cells in 24-well plates were transfected with 500 ng of the psiCHECK-2 reporter plasmids mentioned above (G4Q27, G4M27, and native psiCHECK-2) using Lipofectamine 3000 (Invitrogen, ThermoFisher Scientific) in accordance with the manufacturer’s instructions. The ratios of luciferase activity for the renilla and firefly luciferases were measured 48 hrs (hours) after transfection using the dual-Luciferase reporter assay kit from Promega and a luminometer. The ratio of luciferase activity in the G4Q27 and G4M27 reporter constructs was normalized to the values determined for the empty psiCHECK-2 vector.

### 2.10 Quantitative mRNA Expression

In continuation of our study, we are interested to see whether this PRCC G-quadruplex motif inhibits expression of the *PRCC-TFE3* fusion gene at the transcriptional level or not. For this study, UOK146 cells were treated with 5µM of PDS for 24 hours. Additionally, cells were transfected with 200 nM of siRNA against the *PRCC-TFE3* fusion junction (5’- AAGGGAGATGTGCAGACCCAT-3’) and universal negative control siRNA as a negative and positive control, respectively, using Lipofectamine 3000 as per the manufacturer’s protocol. Total cellular RNA was extracted from PDS-treated and siRNA transfected cells using TRIzol reagent followed by DNaseI treatment to eradicate any remnants of DNA contamination. Then, using a cDNA synthesis kit (Applied Biosystems, Waltham, MA) in accordance with the manufacturer’s instructions, cDNA was produced from 1 µg of pure RNA. Thermo Scientific’s SYBR green Real-time PCR master mix was used to perform qRT-PCR (Quantstudio 6, ThermoFisher Scientific) in triplicates to determine the fusion gene expression levels using *PRCC-TFE3* fusion gene primers (FP:5’-CCAAGCCAAAGAAGAGGAAA-3’ and RP:5’- CGAGTGTGGTGGACAGGTACT-3’).

### 2.11 Cell culture and Immunofluorescence

HEK293T cells were grown on methanol-charged coverslips in the 6 well plates. The next day, cells were transfected with 1 μg of the fusion plasmid construct. The next day, the media were replaced. 48 hrs after post-transfection, cells were treated with 3µM of PDS and 100 mM KCl for another 10–12 hrs to induce/stabilize G-quadruplex structure in genes inside the cells followed by washing with PBS and fixed with 3.7% paraformaldehyde for 15 min, followed by PBS washing. Alternatively, for categorizing both RNA and DNA G-quadruplex formations in the fusion gene construct, cells were incubated with Dnase and RNase, respectively, for 1 hr at 37°C after the fixation. Cells were permeabilized with 0.1% Triton X-100 for 30 min at room temperature, followed by washing with PBS. Blocking was done with 4% bovine serum albumin for 2 hrs at room temperature. After blocking, cells were incubated with G-quadruplex-specific antibody BG4 at 1:100 dilutions for 3 hrs at room temperature in a moist condition, followed by washing with PBS and incubation with the primary antibody (Flag) at 1:500 dilutions for 3 hrs at room temperature. After incubation with Flag antibody, cells were washed with PBS and incubated for 3 hrs at room temperature with secondary antibody Alexa Fluor-546 in 1:500 dilutions. Cells were washed with PBS, and nuclear staining was done with 1 μg/ml DAPI for 15 min, followed by washing and air drying. Slides were mounted using DPX and sealed with nail polish. Images were captured under a confocal microscope (ZWISS LSM 780). Image analysis or fluorescent intensity measurement was done by imageJ software.

## 3. Results

### 3.1 Circular Dichroism Study

CD spectroscopy was done to check the formation of the G-quadruplex conformation in the target sequence. CD spectra of PRCC G4Q show a negative peak at 240 nm and a positive peak around 263 nm (Figure 1), supporting the parallel type G-quadruplex structure [17]. In contrast, the CD spectra of the PRCC G4M sequence show a different peak from G-quadruplex structure even in the presence of 100 mM KCl, confirming non-G-quadruplex structure. As a positive control, the human telomeric sequence (TelQ) was also utilized. With a negative peak at 240 nm, a shoulder peak at 265 nm, and a positive peak at 290 nm, the CD spectra of human telomere G-quadruplex DNA showed a mixed hybrid type. Following that, CD spectra of PRCC G4Q were acquired in the presence of the well-known G-quadruplex stabilizer PDS. The G4Q oligo sample exhibits a negative peak at 240 nm and a positive peak around 263 nm in the presence of PDS, which are traits of the parallel quadruplex structure; however, the peak intensity around 263 nm and 240 nm get reduced in the presence of PDS with no change in peak position compared to without PDS, the G4Q only oligo sample (Figure 2). These results confirm the formation of a G-quadruplex structure within the *PRCC-TFE3* fusion gene around the *PRCC* gene segment.

**Figure 1:**
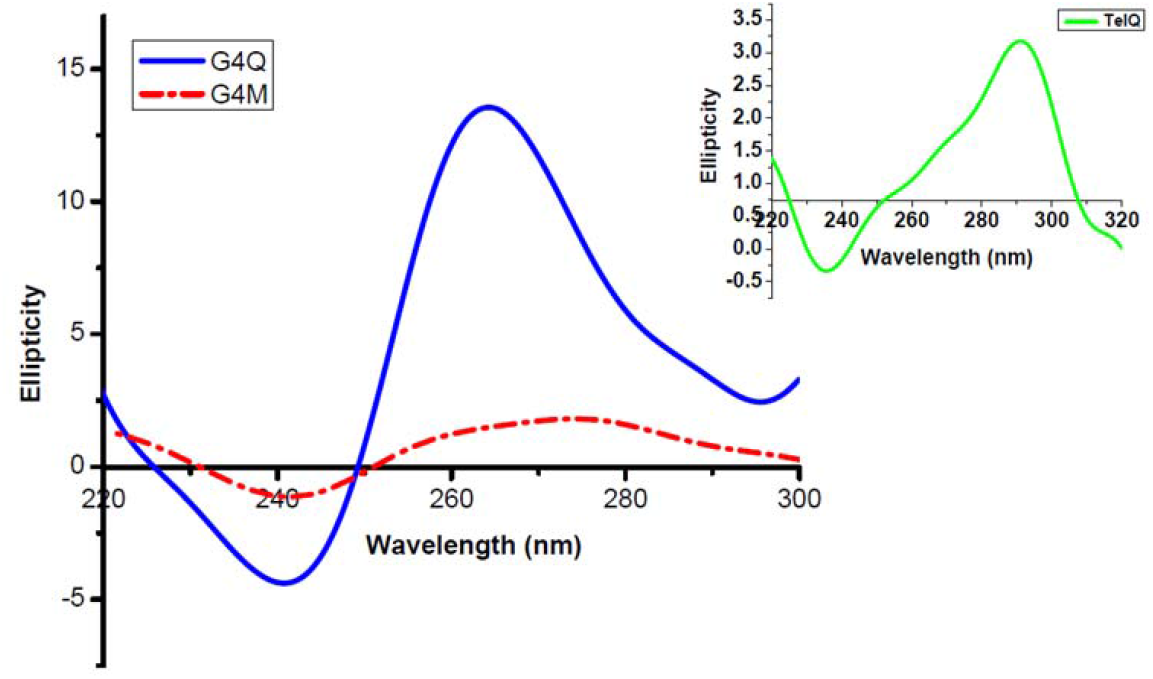
CD Spectra of G4Q (G-rich sequence within the PRCC stretch of PRCC-TFE3 fusion gene) and G4M (mutated form of G4Q, where G is replaced by A) and human telomeric (TelQ) sequence (Inset). Showing formation of parallel G-quadruplex structure in G4Q and mixed hybrid G-quadruplex in TelQ while spectra of the PRCC G4M sequence shows non G- quadruplex structure

**Figure 2:**
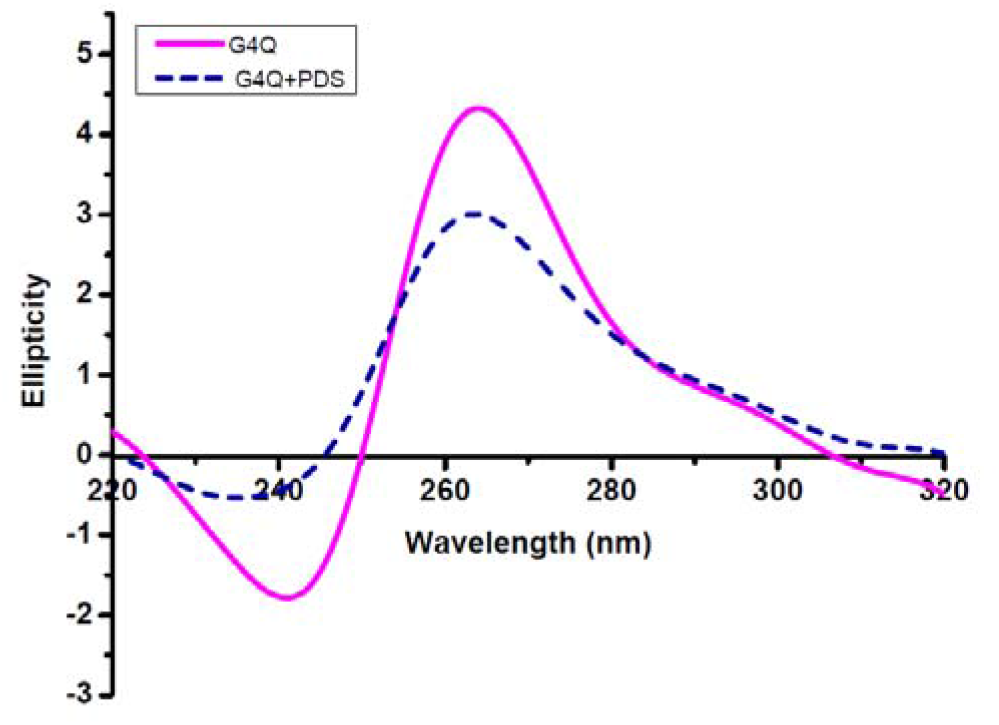
CD spectra of PRCC G4Q in the absence or presence of 5μM pyridostatin (PDS). Showing formation of parallel G-quadruplex structure in G4Q in the presence of PDS. The peak intensity around 263 nm and 240 nm get reduced in the presence of PDS

### 3.2 Thermal Stabilization of G-quadruplex

Thermal stabilization of *PRCC* G4Q G-quadruplexes in the presence of PDS was performed using CD melting at 295 nm. In both cases, with and without PDS, the melting temperatures (Tm) of each transition were calculated and compared by plotting the fraction-folded versus temperature graph. The fraction folded was calculated using the following formula: Q_t_-Q_u_/Q_f_-Q_u_, where Q_t_ is the molar ellipticity at temperature T and Q_u_ and Q_f_ are the molar ellipticities of the unfolded (highest temperature) and completely folded (lowest temperature), respectively. We observed that the Tm of PRCC G4Q was 63.2°C which increases up 69°C with the addition of 5 µM PDS. Thus, 5 µM of PDS increases the stability of *PRCC* G4Q G-quadruplex by approximately 6°C (Figure 3). This finding indicates that the G-quadruplex specific ligand stabilized the newly discovered G-quadruplex.

**Figure 3:**
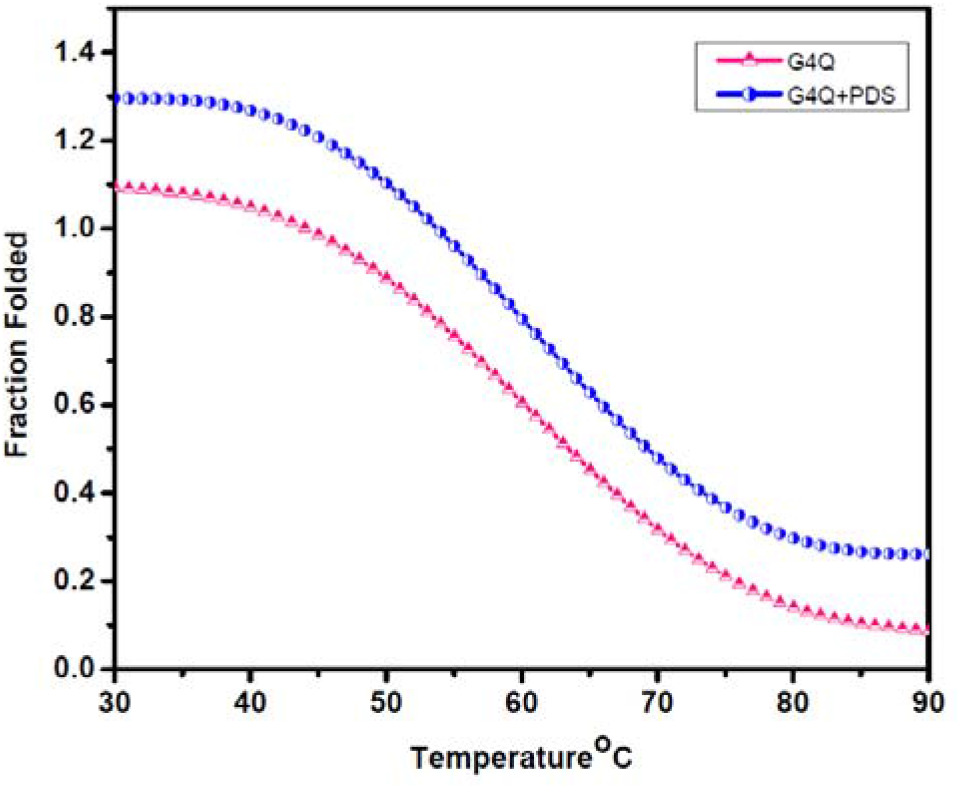
CD melting curve of 5 μM PRCC G4Q in the absence or presence of 5μM PDS. Showing increment in melting temperature (Tm) of G4Q in the presence of PDS by 6°C

### 3.3 UV-Vis Spectroscopy

UV-Vis absorption spectroscopy was used to investigate the interaction of PDS with *PRCC* G-quadruplex (G4Q), as shown in Figure 4. PDS showed two absorption peaks around 310 and 325 nm. With the addition of increasing concentrations of G4Q oligo from 0.5 to 10 μM, absorption peak of PDS decreased around 310 nm, suggesting the formation of PDS-G-quadruplexes or interaction between PDS and G4Q. Simultaneously, no broad change in peak intensity was found with the addition of G4M samples (Figure 4). These results also support the formation and stabilization of G-quadruplex conformation in the targeted sequence, as PDS is known as a binder of G-quadruplex conformation rather than duplex or non G-quadruplex conformation.

**Figure 4:**
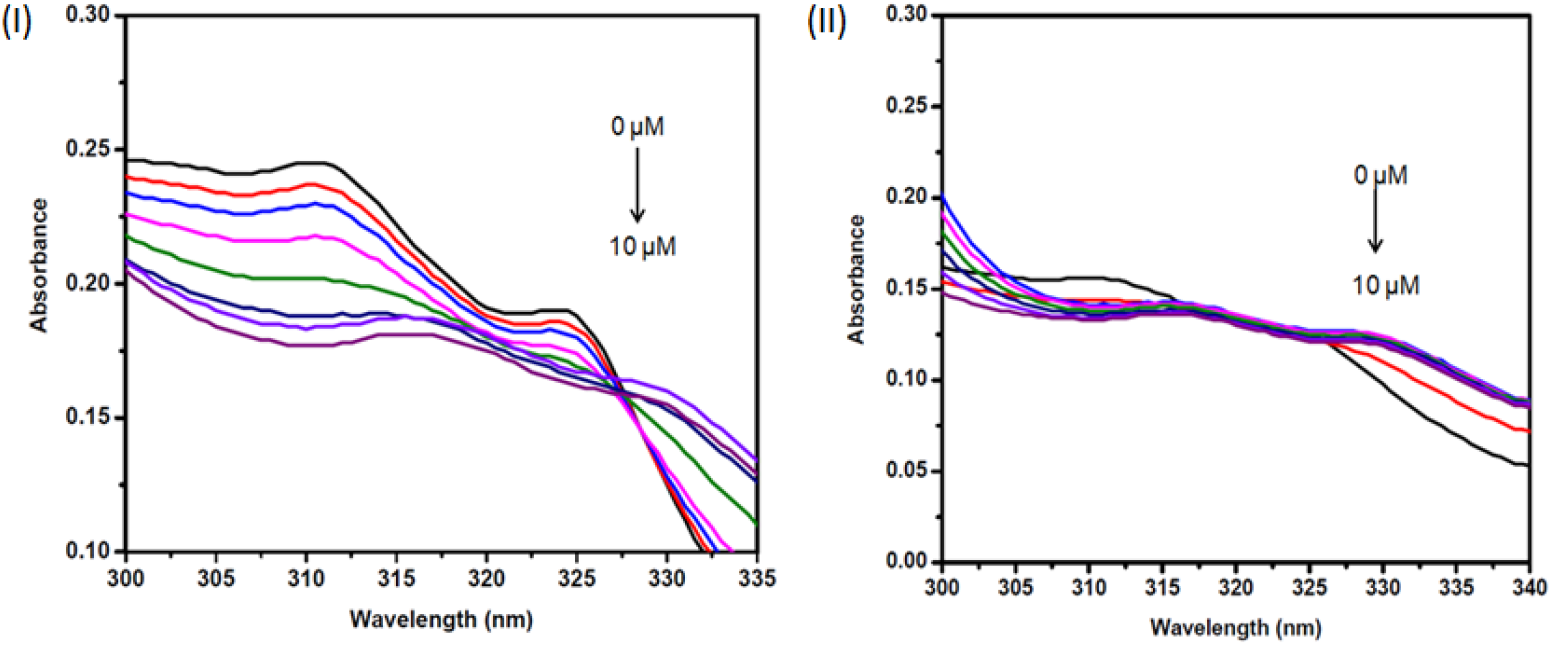
Absorption spectra of PDS (5 μM) in the absence and presence of increasing concentration (0, 0.5,1, 2, 4, 6, 8, and 10 μM) of PRCC G4Q (I) and PRCC G4M (II). Showing two absorption peak at 310 & 324 nm which get reduced by increasing concentration of PRCC G4Q. while no broad change was observed by increasing concentration of PRCC G4M

### 3.4 Binding Parameters by Fluorescence Spectroscopy

Fluorescence quenching in PDS was observed during the interaction of G-quadruplex (*PRCC* G4Q) with PDS. The fluorescence spectrum of PDS displayed its highest peak at 537.5 nm. PDS exhibited emission in the range of 520–575 nm. A decrease in the fluorescence intensity at 537.5 nm with increasing concentration of G4Q (0.5-10 μM) indicates its binding to the G-quadruplex structure (Figure 5). The fall in PDS’s fluorescence intensity suggests that the substance’s ability to bind to the G-quadruplex has effectively declined in quantum efficiency. The fluorescence data were analyzed using the Stern-Volmer equation (Eq.1) [18] and double log plots of F_0_-F/F versus [Q] were created.

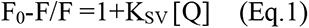

where Ksv is the Stern Volmer constant, Q is the ligand concentration, and F and F_0_ are the fluorescence intensities in the presence and absence of the quencher (G4Q), respectively. The modified Stern-Volmer equation (Eq.2) was used to assess binding affinity and the number of binding sites.

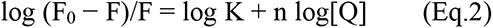

**Figure 5:**
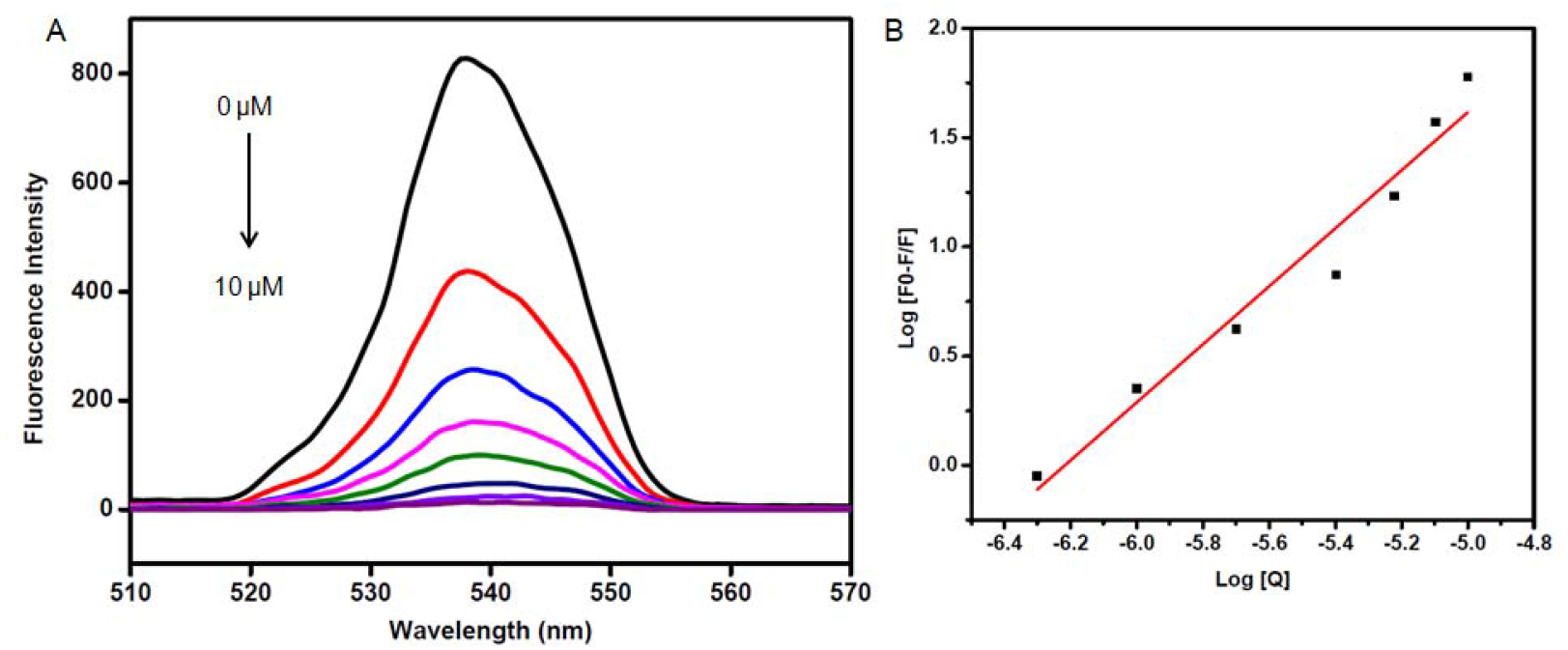
(A) Fluorescence emission spectra of PDS (5 μM) in the absence and in the presence of increasing concentration of PRCC G4Q (0, 0.5, 1, 2, 4, 6, 8, and 10 μM). Fluorescence quenching at 537.5 nm with increasing concentration of G4Q suggests substance’s ability to bind to the PRCC G4Q. (B) The double logarithmic plot to calculate binding constant and binding number

Where K and n are the corresponding binding constant and number of binding site. The Stern Volmer constant (Ksv) for the *PRCC* G4Q-PDS interaction was found to be 2.073×10^6^M^-1^ and the binding constant (K) was found to be 8.25×10^6^M^-1^ and number of binding site was 1. The values of K and n for the G quadruplex-PDS interaction were calculated from the intercept and slope of the plot of log (F_0_- F)/F vs. log [Q] (Figure 5).

### 3.5 PCR Stop Assay

The *PRCC* G4Q oligo produced stable G-quadruplex structures with increasing concentrations of PDS (0, 2, 3, 5, 7, and 10 μM) and prevented the formation of a double-stranded PCR product a the band of the PCR product became faint and disappeared with increasing concentrations of PDS (Figure 6), indicating that the G-quadruplex structure hampered polymerase extension.

**Figure 6:**
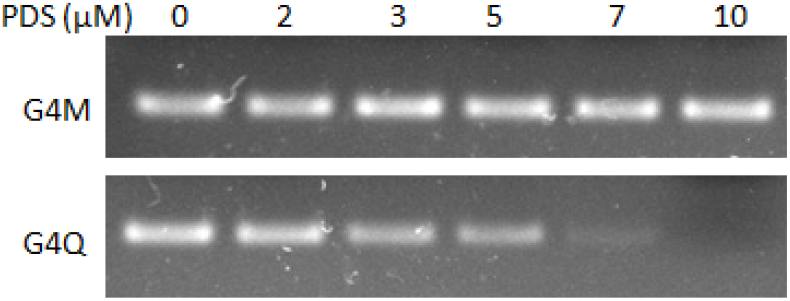
Gel picture showing the Taq polymerase extension inhibition leading to the PCR stop at 0, 2, 3, 5, 7, and 10 μM PDS in PRCC G4Q and G4M. G-quadruplex stabilization by PDS prevents oligomer annealing and subsequently Taq polymerase extension. In PRCC G4Q, PCR inhibition increases with increasing concentration of PDS, while extension is unaffected by mutant type oligo G4M

Whereas in case of *PRCC* G4M oligo, PCR product’s band not getting disappear with increasing concentration of PDS that means polymerase extension was not inhibited as *PRCC* G4M oligo is unable to form a stable G-quadruplex because every G stretch in this oligo has been altered by adenine. This assay demonstrated the formation and stabilization of a G-quadruplex structure within the *PRCC* stretch of the *PRCC-TFE3* fusion gene, as well as the inhibition of polymerase extension activity by this G-quadruplex motif activity.

### 3.6 Transcription Inhibition of Fusion Gene by G-quadruplex

The expression fold of the *PRCC-TFE3* fusion transcript in the UOK 146 cell line is reduced by 0.54 in PDS-treated cells, while it is reduced by 0.20 in the positive control, i.e., cells with the fusion transcript knocked down by siRNA (Figure 7). As a control, a universal negative control siRNA was transfected into UOK 146 cells. The qRT-PCR experiment confirmed the repression of the *PRCC-TFE3* fusion gene as a result of the G-quadruplex motif within the transcript, as PDS has already been shown to stabilize the G-quadruplex structure in the PRCC stretch of the *PRCC-TFE3* fusion gene by thermal melting study.

**Figure 7:**
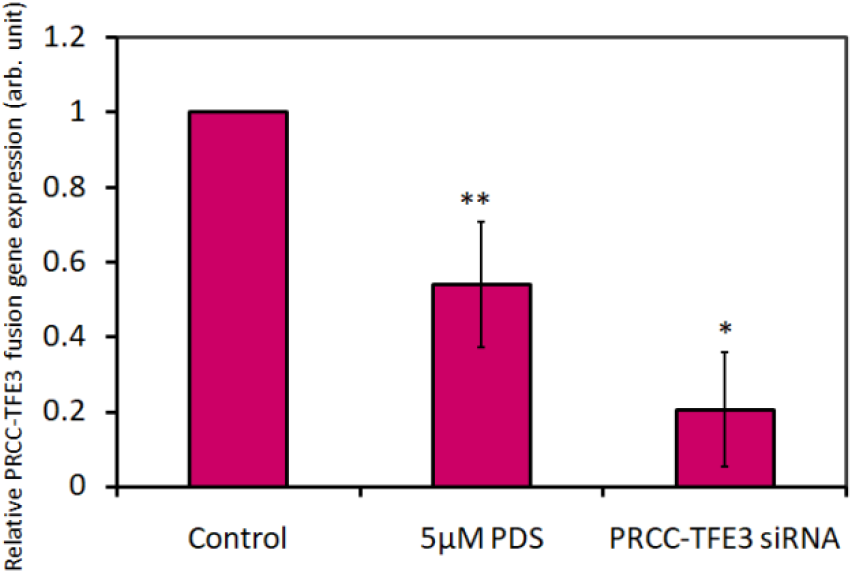
Relative fold change of *PRCC-TFE3* fusion oncogene in UOK146 cell line by stabilizing G-quadruplex structure using 5 μM PDS. Expression fold of the fusion transcript is reduced by 0.54 in PDS-treated cells, and 0.20 in fusion transcript knocked out cells, suggesting the repression of the *PRCC-TFE3* fusion gene in the effect of G-quadruplex stabilizing ligand PDS.

### 3.7 Cellular Visualization of G-quadruplex

As shown in Figure 6, the signal of BG4 or G-quadruplex foci was found to be enhanced in fusion clone-transfected HEK293T cells in comparison to untransfected cells. Both inside the nucleus and around the cytoplasm, BG4 signal or G-quadruplex foci were seen, which correspond to DNA and RNA G-quadruplex, respectively [19–20], and both of them are significantly increased in fusion gene transfected cells. The mean fluorescence intensity of endogenous BG4 signals in untransfected cells was 5.7, which gets enhanced up to 15.3 in transfected cells (Figure 8). Further DNase treatment results in the loss of the BG4 signal inside the nucleus in fusion clone transfected and PDS treated cells, whereas in RNase treated cells, signals are lost from the cytoplasm, corresponding to losses in DNA and RNA G-quadruplexes, respectively (Figure 9).

**Figure 8:**
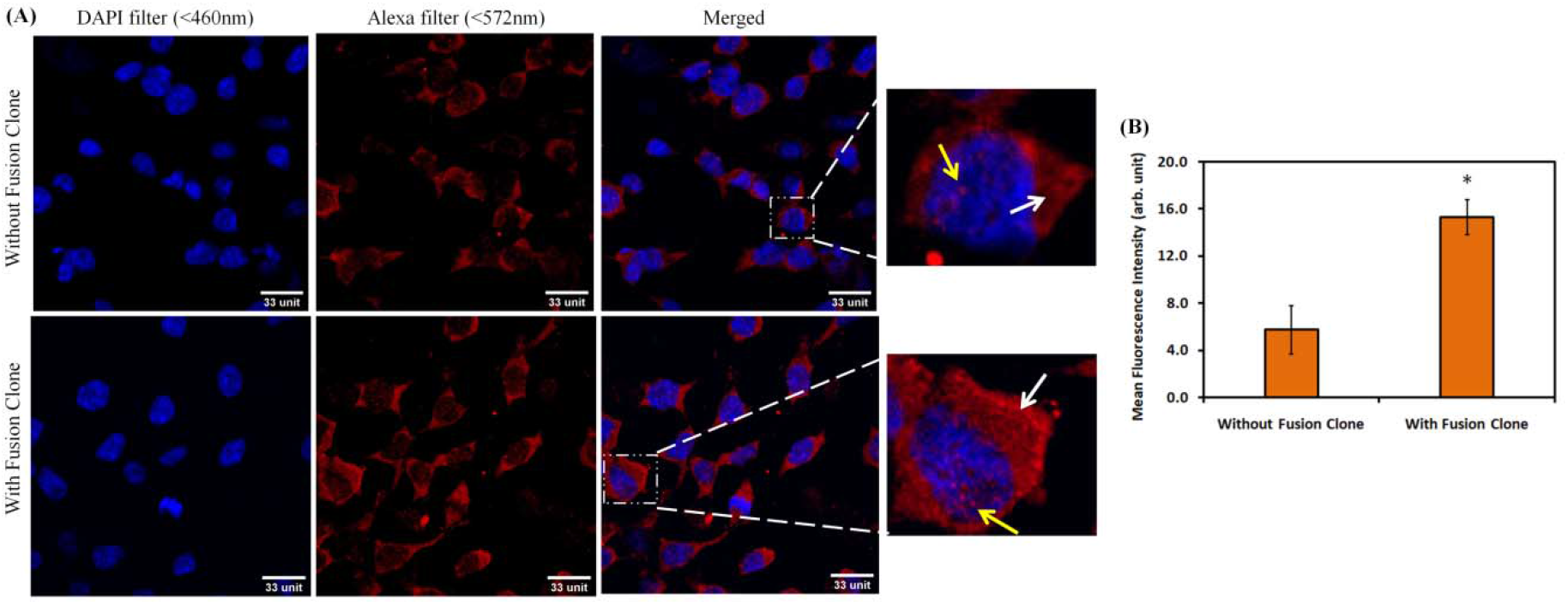
(A) Confocal imaging of G-quadruplex with or without fusion clone transfected HEK293T treated with 3μM PDS and 100 mM KCl to stabilize G-quadruplex structure in the cells. G-quadruplex foci (red) has been enhanced in cells with clone, suggests increment in G-quadruplex formation compared to without clone. White & yellow arrows in insets indicate representative clusters of RNA & DNA G-quadruplex respectively. Nuclei were counter stained by DAPI. Scale bar 33 μm. (B) Bar graph of mean fluorescence intensity of endogenous G-quadruplex foci. Mean fluorescence intensity in untransfected cells was 5.7, which gets enhanced up to 15.3 in transfected cells. Error bars represent the S.D. of three independent experiments. *P ≤0.05, without clone vs. with clone

**Figure 9:**
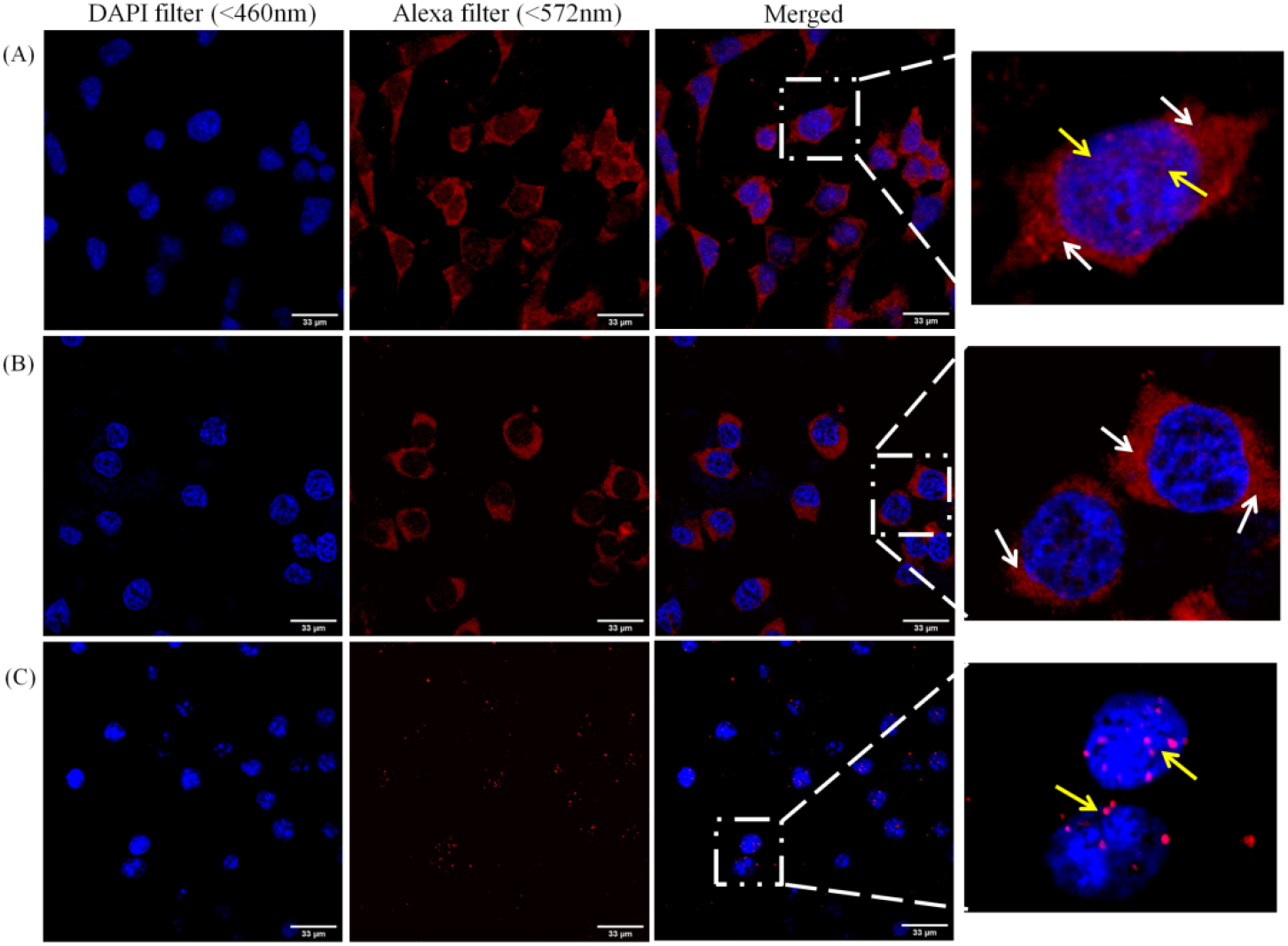
Confocal imaging of G-quadruplex in fusion clone transfected HEK293T cells treated with 3 μM PDS & 100 mM KCl to stabilizes the G-quadruplex structure inside the cells. White & yellow arrows in insets indicate representative clusters of RNA & DNA G-quadruplex respectively. Nuclei were counter stained by DAPI. (A) Cells treated with PDS only: showing both DNA & RNA G-quadruplex. (B) Cells treated with PDS+Dnase: only RNA G-quadruplex (cytoplasmic signal). (c) Cells treated with PDS+Rnase: only DNA G-quadruplex (nuclear signal). Scale bar 33 μm.

### 3.8 Translation Inhibition by G-quadruplex in Living Cells

Next, we were interested in investigating the impact of the G-quadruplex motif (G4Q motif) upon translation in living eukaryotic cells. Relative luciferase activity in the G4Q27 and G4M27 constructs was found to be 60.9% and 90.2%, respectively. Figure 10 depicts graphs of relative luciferase activity. Surprisingly, 40.5% less protein is synthesized when the G4Q27 motif is inserted upstream of the renilla luciferase start codon. However, translation is not greatly impacted by the mutant form G4M27, which does not form a stable quadruplex structure. The G-quadruplex’s partial suppression of translation falls within the range of previous G-quadruplexes’ observations [21]. This observation is consistent with the hypothesis that the *PRCC-TFE3* fusion gene develops a G-quadruplex structure that prevents translation. According to what we’ve seen, this G-quadruplex motif can grow inside of cells and can stop reporter translation.

**Figure 10:**
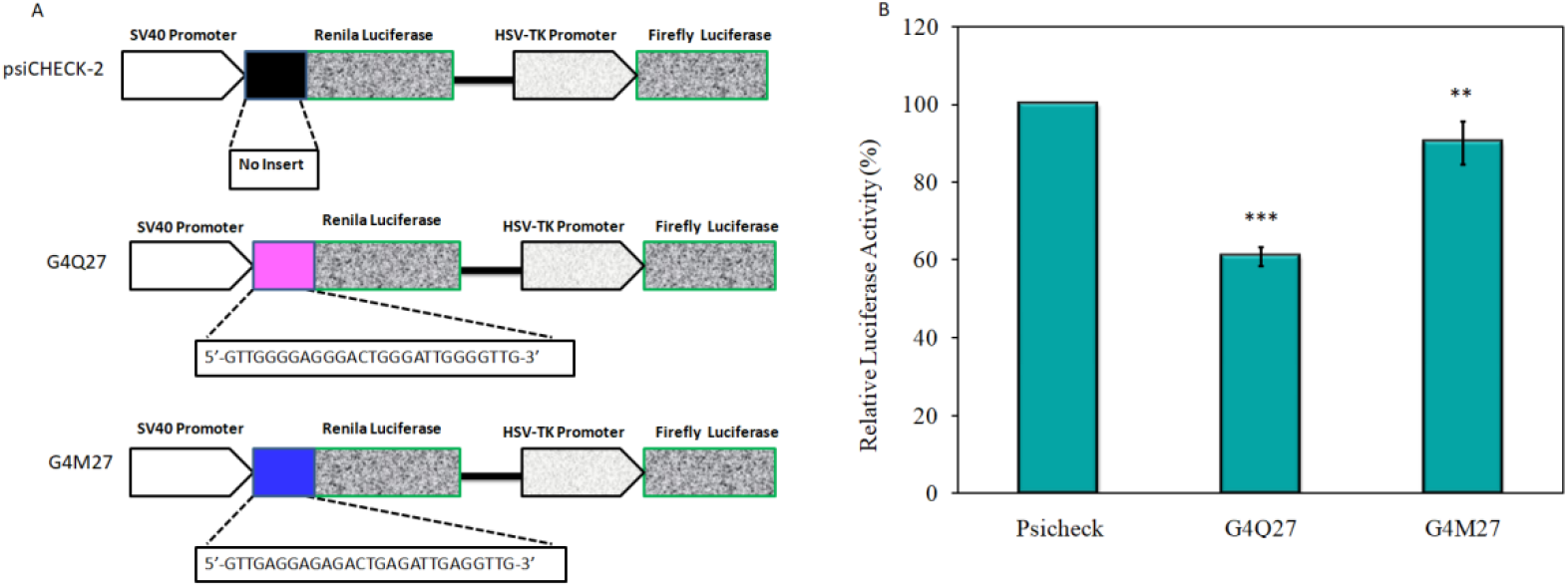
(A) Schematic representation of construction of plasmid. G4Q27 (G-rich sequence within the PRCC stretch of *PRCC-TFE3* fusion gene) and G4M27 (mutated form of PRCC G4Q). psiCHECK-2 is empty vector without any change. (B) Relative translation efficiency of the three constructs measured by quantifying luciferase activity. Data were normalized by empty vector psiCHECK-2. G-quadruplex’s partial suppression of translation has been seen as, the relative expression of the renilla gene in G4Q27 get significantly reduced to 40% while only 10% reduction has been found in G4M27. Error bars represent the S.D. of three independent experiments. *P ≤0.05, **P ≤ 0.01, ***P ≤ 0.001, control vs. other groups

## 4. Discussion

G-quadruplex structures are viewed as new molecular targets for cancer therapies due to their direct visibility in living cells [22]. Putative G-quadruplex forming sequences or G-quadruplex structures in oncogenes provide new opportunities for developing anticancer tactics [16]. In this study, we are primarily interested in the *PRCC-TFE3* fusion gene, which is thought to be an oncogene in Xp11.2 translocation papillary renal cell carcinoma with increased transcriptional activity [4, 13]. We examined the *PRCC-TFE3* fusion gene sequence using online web servers like non BdB and QGRS mapper, and we discovered the stretches of guanine residues that had a propensity to generate G-quadruplex secondary structures. Preliminary sequence analysis revealed that this *PRCC-TFE3* fusion gene contains numerous PQS sequences. Of these PQS, the one with the highest score is located in the exonic region of the *PRCC* stretch, as reported in our published review paper [13]. Therefore, in the current study, we selected this sequence stretch of fusion oncogene for *In vitro*, biophysical/biochemical, and cellular analysis to determine the G-quadruplex formation and impact on the expression machinery of this fusion oncogene. The PQS of the exonic *PRCC* section of the *PRCC-TFE3* fusion gene was assumed to be extremely important for the study since it passes on to the fusion transcript as well as the fusion oncoprotein coding region also upstream to the translation start site. The structure of folded nucleic acid G-quadruplexes has been extensively studied using circular dichroism (CD) spectroscopy. Also, to distinguish the parallel, anti-parallel, and mixed-type secondary structures of G-quadruplex DNA, CD is a frequently used technique. Its extreme sensitivity to even minute changes in the mutual orientation of DNA’s absorbing bases makes CD one of the most used techniques for analyzing the conformational characteristics of nucleic acids, particularly those of guanine quadruplexes [23]. By CD spectra, we have shown the formation of parallel G-quadruplex structure in G4Q oligo, a brief stretch inside the *PRCC-TFE3* fusion gene, while its mutated form, G4M oligo, fails to achieve G-quadruplex structure. Further CD spectra of *PRCC* G4Q in the presence of the renowned G-quadruplex binder PDS were also recorded to see the effect of PDS on this newly identified G-quadruplex. PDS is a highly selective G-quadruplex DNA-binding small molecule intended to interact with and stabilize G-quadruplex structures [24]. Additionally, PDS is also known to cause telomere disruption and activate a DNA damage response [25]. The *PRCC* G4Q DNA sequence still exhibits parallel G-quadruplex conformation in the presence of PDS, but intensity peaks around 263 nm and 240 nm get reduced in the presence of PDS with no change in peak position, suggesting the formation of G-quadruplex structure in selected PRCC segments of the *PRCC-TFE3* fusion gene. To examine how a ligand interacts with and affects the stability of quadruplex DNA, DNA melting experiments have been widely used. The midway point of a certain transition, where the complex is 50% dissociated, is known as the melting temperature (Tm). The Tm value only provides a rough assessment of the prevailing shape. At temperatures above the Tm, the structure is primarily unfolded, whereas at temperatures below the Tm, the structure is primarily folded [26]. With the addition of 5 μM of PDS, the Tm of a targeted PRCC G4Q G-quadruplex is increased by 6°C, an indication of the stabilization of *PRCC* G4Q by PDS. A ligand’s DNA-binding behavior is routinely studied by UV-visible absorption. Therefore, we looked into how PDS interacted with the *PRCC* G4Q G-quadruplex by UV-visible titration of increasing amounts of G4Q oligo with a fixed concentration of PDS, which showed maximum absorption at 310 nm. The addition of increasing concentrations of G4Q to PDS resulted in a significant decrease in PDS peak absorption at 310 nm, with approximately 25-27% hypochromicities. For comparison, we also checked the interaction between PDS and G4M, and as expected, no drastic change has been found in PDS’s absorption peak at 310 nm. Upon the addition of G4M, smaller hypochromic changes (12–14%) were observed, suggesting interaction of the G-quadruplex binding ligand PDS with *PRCC* G4Q. Further, in order to determine the binding parameters in the PDS-PRCC G4Q interaction, steady state fluorescence titration was carried out as part of the PDS-G-quadruplex binding investigation. We examine the possibility that a compound’s interaction with the G-quadruplex would change its fluorescence characteristics. The steady-state fluorescence spectrum shows the contribution of each emissive species in solution, making it possible to track changes in the chromophore’s surroundings. The fluorescence spectra of PDS shows a single peak around 537- 540 nm. In this analysis, we found strong fluorescence quenching in the PDS’s fluorescence spectrum with the addition of an increasing amount of *PRCC* G4Q G-quadruplex oligo. Fluorescence quenching is the reduction of a fluorophore’s quantum yield of fluorescence brought on by a variety of molecular interactions with the quencher molecule, including excited-state reactions, molecule rearrangements, energy transfer, the formation of ground-state complexes, and collisional quenching [27]. The modified Stern-Volmer equation was also used to calculate the binding constant and number of binding sites in the PDS-G4Q binding parameters. These all biophysical techniques together support the presence or formation of the G-quadruplex conformation within the targeted stretch of *PRCC-TFE3* fusion oncogene. Next, we are interested in seeing the formation and existence of G-quadruplex in this selected gene in cellular conditions as well as its impact upon their expression machinery. DNA gets unwinds during the transcription and replication processes, and PQS can fold into a G-quadruplex structure prior to reannealing, which therefore influences those processes [26]. A PCR stop assay was used to investigate the effect of PRCC G-quadruples structure stabilization on replication blockage. The PCR stop assay is a promising assay for investigating the effect of polymerase extension activity on G-quadruplex structure. Primer extension by Taq polymerase was halted as a result of the stabilization of the G-quadruplex structure by the ligand inside the template sequence [28]. The *PRCC* G4Q DNA oligomer stabilizes into a G-quadruplex structure in the presence of PDS, which blocks further processing of Taq DNA polymerase and results in loss of amplification or double-stranded PCR product. In the case of G4M oligo, guanine stretches were mutated by adenine, which reduced polymerase extension blocking because no G-quadruplex structure was present to obstruct polymerase activity. These findings demonstrate the potential threat posed by a stable G-quadruplex structure to the replication of the target fusion oncogene. Next, we checked the impact of G-quadruplex formation upon transcription machinery. Several *In vitro* experiments suggest that the transcribed region’s G-quadruplex motif can obstruct the movement of the RNA polymerase complex. The co-transcriptional formation of intramolecular G-quadruplex DNA on the template strand as well as the intermolecular G-quadruplex DNA made up of the non-template DNA strand and the nascent RNA would occur when the G-quadruplex motifs are present on either strand within the transcribed region and would physically obstruct subsequent rounds of the RNA polymerase’s movement [29]. UOK 146, a cell line carrying the *PRCC-TFE3* fusion gene, was used for this study. Cells treated with PDS showed a significant reduction (53%) in fusion gene transcripts. PDS is a G-quadruplex stabilizing agent, and when PQS is treated with it, it forms stable G-quadruplexes. These stable structures pose a major threat to RNA polymerase and transcription [13]. These findings suggest that the alteration in *PRCC- TFE3* fusion transcript levels can be attributed to interference with the transcriptional machinery caused by G-quadruplexes. Previous research has demonstrated that G4Q motifs in the 5’-UTR of the mRNA can inhibit translation, viz. NRAS [16] and human Zic-1 zinc-finger protein [30]. In order to clarify this essential factor, we carried out dual luciferase assays. The effect of G-quadruplex on translation was investigated using *PRCC* and its mutant minigene dual reporter constructs, in which the G-quadruplex motif was located immediately upstream of the renilla gene while the firefly gene remained unmodified, to determine whether the presence of a G-quadruplex structure right upstream of the start site may affect a gene’s expression. Therefore, as a normalizing control, the firefly expression was used. In comparison to the empty vector, the relative expression of the renilla reporter in the G4Q27 construct was significantly reduced to about 40%. The G-quadruplex’s partial suppression of translation falls within the range of other G-quadruplexes that have previously been observed [22, 31]. Additionally, only a 10% drop was seen in the construct with the altered G-quadruplex motif at G4M27, which indicated a remarkable recovery to luminescence levels comparable to the empty vector. The G4M27 sequences, as previously stated, do not exhibit any CD signature of G-quadruplex conformation, which is worth mentioning at this point. This recovery of luminescence caused by G-quadruplex disruption makes a substantial contribution to the body of research showing that the effect was mainly caused by the G-quadruplex motif. This suggests that the efficiency of translation is reduced when G-quadruplex is present after the translation start site. These results support the hypothesis that a G-quadruplex structure is formed in the *PRCC* stretch of the *PRCC-TFE3* fusion gene, which can inhibit the replication and translation of the fusion gene or transcript. Encouraged by the *In vitro* results, we were therefore eager to use confocal microscopy to observe the G-quadruplex in the target gene under cellular conditions. In order to do this, we transfect the HEK293T cells with the full-length *PRCC-TFE3* fusion plasmid and allow them to develop for roughly 48 hours so that the level of the fusion gene in the cells is within the range of detection. Following this, 3 μM PDS and 100 mM KCl were administered to both transfected and non-transfected cells to stabilize the G-quadruplex inside the cells, as a gene’s ability to form G-quadruplexes is regulated by the course of the cell cycle and endogenous G-quadruplex structures can be stabilized by a small-molecule ligand or G-quadruplex stabilizer. It has also been reported a remarkable increment in BG4 signal in PDS-treated cells as compared to non- treated cells [19]. We therefore assumed that PDS treatment of cells would result in effective G-quadruplex targets by the BG4 antibody. Fluorescence intensity was measured and compared in order to monitor the G-quadruplex development within the cells. G-quadruplex-specific antibody BG4, which is able to identify both DNA [19] and RNA [20] G-quadruplex in human cells, was employed for this G-quadruplex-specific assay. Since the BG4 antibody has a FLAG tag sequence, it is further detected by a FLAG antibody using an anti-Flag Alexa Fluor 546 secondary antibody. In transfected cells, the mean fluorescence intensity of G-quadruplex- bounded antibodies was found to be 9.6 times higher than in non-transfected cells. Human cells contain millions of PQS-containing genes, including telomerase, and these PQS can fold into G-quadruplex structure during the S-phase or DNA replication phase [19], which could explain why signals were observed in untransfected cells. The existence of the *PRCC-TFE3* fusion gene, which also carries the PQS, may be the reason for the increase in G-quadruplex foci in transfected cells. Following a breakthrough in the cellular visualization of fusion gene G-quadruplexes, we were eager to learn whether both DNA and RNA G-quadruplexes could form inside the cell in this fusion gene. For this, fusion plasmid-transfected and PDS-treated cells were exposed to DNase and RNase treatment for a selective visualization of RNA and DNA G-quadruplexes, respectively. When cells were treated with RNase, the initially enhanced G-quadruplex signal in transfected cells was clearly eliminated from the cytoplasmic area, indicating the absence of an RNA G-quadruplex signal. Similarly, in DNase-treated cells, the signal of G-quadruplex foci inside the nucleus disappears, indicating the loss of DNA G-quadruplex structure. Taken together, these results support the hypothesis of the formation of a G-quadruplex structure inside the *PRCC* stretch of the *PRCC-TFE3* fusion oncogene, which is also sustained in cellular conditions at both the DNA and RNA levels, and whose stabilization may be capable of controlling the fusion gene expression at both the replication and translation levels.

## 5. Conclusion

Finally, we discovered the possibility of G-quadruplex secondary structure formation upstream of the translation start site inside the *PRCC* segment of the *PRCC-TFE3* fusion gene, an oncogene in translocation papillary renal cancer. Our *In vitro* study demonstrates the formation as well as stabilization of the G-quadruplex structure in the targeted sequence in the presence of the G-quadruplex stabilizing ligand PDS inside the cell. Additionally, stabilization of the G-quadruplex structure in the target gene showed the capacity to inhibit target gene replication, translation, and transcription, suggesting a potential therapeutic strategy against fusion gene-mediated translocation papillary renal cancer. However, since the chosen gene sequence is present in the wild-type *PRCC* as well, additional research is still required to specifically target this secondary structure in the *PRCC-TFE3* fusion gene.

## Acknowledgements

We acknowledge Central Discovery Centre (CDC), Banaras Hindu University, Varanasi for Circular Dichroism facility. We acknowledge ISLS, Banaras Hindu University for providing facility for confocal microscopy. We acknowledge Dr. D.S Pandey, Professor, Department of Chemistry, Banaras Hindu University for providing Spectroscopy facility. We acknowledge W M Linehan, National Cancer Institute, USA for providing the UOK 146 cell line.

## Author Contributions

Neha designed and performed the experiments, and wrote the manuscript. P.D. designed the experiment, review and supervision.

